# Systematic identification of cancer-specific MHC-binding peptides with RAVEN

**DOI:** 10.1101/193276

**Authors:** Michaela C. Baldauf, Julia S. Gerke, Andreas Kirschner, Franziska Blaeschke, Manuel Effenberger, Kilian Schober, Rebeca Alba Rubio, Takayuki Kanaseki, Merve M. Kiran, Marlene Dallmayer, Julian Musa, Nurset Akpolat, Ayse N. Akatli, Fernando C. Rosman, Özlem Özen, Shintaro Sugita, Tadashi Hasegawa, Haruhiko Sugimura, Daniel Baumhoer, Maximilian M. L. Knott, Giuseppina Sannino, Aruna Marchetto, Jing Li, Dirk H. Busch, Tobias Feuchtinger, Shunya Ohmura, Martin F. Orth, Uwe Thiel, Thomas Kirchner, Thomas G. P. Grünewald

**Author notes:** Corresponding author: Thomas G. P. Grünewald, MD, PhD Max-Eder Research Group for Pediatric Sarcoma Biology, Institute of Pathology, Medical Faculty, LMU Munich, Thalkirchner Str. 36, 80337 Munich, Germany, Web: www.lmu.de/sarkombiologie. These authors share first authorship (Michaela C. Baldauf, Julia S. Gerke).

## Abstract

Immunotherapy can revolutionize anti-cancer therapy if specific targets are available. Recurrent somatic mutations in the exome can create highly specific neo-antigens. However, especially pediatric cancers are oligo-mutated and hardly exhibit recurrent neo-antigens. Yet, immunogenic peptides encoded by cancer-specific genes (CSGs), which are virtually not expressed in normal tissues, may enable a targeted immunotherapy of such cancers. Here, we describe an algorithm and provide a user-friendly software named RAVEN (Rich Analysis of Variable gene Expressions in Numerous tissues), which automatizes the systematic and fast identification of CSG-encoded peptides highly affine to Major Histocompatibility Complexes (MHC) starting from publicly available gene expression data. We applied RAVEN to a dataset assembled from more than 2,700 simultaneously normalized gene expression microarrays comprising 50 tumor entities, with a focus on sarcomas and pediatric cancers, and 71 normal tissue types. RAVEN performed a transcriptome-wide scan in each cancer entity for gender-specific CSGs. As a proof-of-concept we identified several established CSGs, but also many novel candidates potentially suitable for targeting multiple cancer types. The specific expression of the most promising CSGs was validated by qRT-PCR in cancer cell lines and by immunohistochemistry in a comprehensive tissue-microarray comprising 412 samples. Subsequently, RAVEN identified likely immunogenic peptides encoded by these CSGs by predicting the affinity to MHCs. Putative highly affine peptides were automatically crosschecked with the UniProt protein-database to exclude sequence identity with abundantly expressed proteins. The predicted affinity of selected peptides was validated in T2-cell peptide-binding assays in which many showed similar kinetics to a very immunogenic influenza control peptide.

Collectively, we provide a comprehensive, exquisitely curated and validated catalogue of cancer-specific and highly MHC-affine peptides across 50 cancer entities. In addition, we developed an intuitive and freely available software to easily apply our algorithm to any gene expression dataset (https://github.com/JSGerke/RAVENsoftware). We anticipate that our peptide libraries and software constitute a rich resource to accelerate the development of novel immunotherapies.

## INTRODUCTION

Immunotherapy is currently transforming clinical oncology and holds promise for cure even for patients with metastatic disease (Mellman et al., 2011; Chen J J Immunol Res 2017). The success of many immunotherapeutic approaches, e.g. adoptive T cell therapy, largely depends on the availability of specific immunogenic target structures presented via Major Histocompatibility Complexes (MHCs) on the surface of cancer cells, but not on that of normal tissues (Coulie et al., 2014). Genetically instable and hyper-mutated cancer entities such as malignant melanoma and lung carcinoma offer such highly specific target structures through missense mutations in the protein coding genome that generate ‘neo-antigens’ (Schumacher and Schreiber, 2015).

However, many cancer types such as pediatric cancers are characterized by a remarkably stable and oligo-mutated genome (Lawrence et al., 2013). In addition, the few recurrent somatic mutations found in pediatric cancers are hardly immunogenic (Orentas et al., 2012). Thus, specific immunotherapy of oligo-mutated cancers is challenging, but may be enabled by the expression of non-mutated cancer-specific genes (CSGs) (Coulie et al., 2014).

Many CSGs are only expressed during early embryogenesis or in immune-privileged germline tissues such as testis (Monk and Holding, 2001; Simpson et al., 2005). This restricted expression pattern increases the likelihood of circulating lymphocytes directed against immunogenic peptides encoded by these CSGs (Simpson et al., 2005), which can be exploited clinically. In neuroblastoma and Ewing sarcoma, which are aggressive and oligo-mutated pediatric cancers (Pugh et al., 2013; Tirode et al., 2014), adoptive T cell therapy targeting CSGs has been successfully applied in humanized mouse models (Blaeschke et al., 2016; Kirschner et al., 2017; Schirmer et al., 2016; Singh et al., 2016) and a first set of patients (Thiel et al., 2017). Screening for additional CSGs could be enabled by comprehensive and already available transcriptome datasets of cancer and normal tissues (Rung and Brazma, 2013), However, due to the lack of specific algorithms and user-friendly tools, the identification of CSGs and derivative peptides with high affinity to MHCs continues to be laborious and slow (Gubin et al., 2015).

To accelerate this process and to identify CSGs suitable for targeting various oligo-mutated cancer entities, we developed an algorithm and provide an intuitive software termed RAVEN (Rich Analysis of Variable gene Expressions in Numerous tissues), which automatizes the systematic and fast identification of cancer-specific peptides with high affinity to MHCs starting from gene expression data. By applying RAVEN to a dataset of more than 2,700 gene expression microarrays comprising 50 tumor entities and 71 normal tissue types, we identified a library of peptides suitable for targeting multiple cancers. Our datasets and software represent a rich resource for the development of immunotherapies.

## MATERIALS AND METHODS

### Microarray data

Publicly available gene expression data generated with Affymetrix HG-U133Plus2.0 microarrays for 3,101 samples comprising 50 tumor entities and 71 normal tissue were retrieved from the Gene Expression Omnibus (GEO) or the Array Express database at the European Bioinformatics Institute (EBI). Accession codes are reported in **Supplementary Table 1**. Microarray quality checks were performed by analyzing the Relative Log Expression (RLE) and Normalized Unscaled Standard Error (NUSE) scores with the Bioconductor packages *affyPLM* (Brettschneider et al., 2008) and *hgu133plus2hsentrezgcdf* (Dai et al., 2005) in the statistical language R (R Development Core Team). The cut-offs for defining high quality were set as (1^st^ quartile – [1.5 x interquartile range]) and (3^rd^ quartile + [1.5 x interquartile range]).

All microarrays were pre-processed (normalized) simultaneously in R with the Robust Multi-chip Average (RMA) algorithm (Irizarry et al., 2003) including background adjustment, quantile normalization and summarization using custom brainarray Chip Description Files (CDF; ENTREZG, v19) yielding one optimized probe-set per gene (Dai et al., 2005).

### Identification of CSG-scores from normalized expression intensities

To identify CSGs in any given gene expression dataset, we calculated the outlier expression of a gene *x* in a specific cancer *c* by considering the adjusted upper quartile mean of its expression signals, as such approach avoids bias through extreme outliers in a tiny subset of samples (above 95^th^ quantile [Q95]) (Xu et al., 2012). The adjusted upper quartile mean, named ‘Outlier Score’ (OS), of gene *x* in cancer type *c* is given as

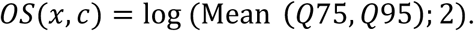

Next, a ‘Penalty Score’ (PS) for gene *x* was calculated on the basis of its adjusted upper quartile mean among different types of normal human tissues *n* a

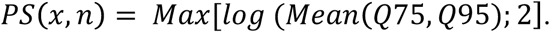

The CSG-score of a gene *x* in a given cancer type *c* was then calculated as

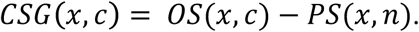

Previously reported algorithms included weighting scores for each normal tissue type based on their possible degree of estimated ‘immuno-privilege’ or even excluded highly immune-privileged organs such as testis from calculation of a PS (Kadota et al., 2006; Xu et al., 2012). In contrast, we considered each normal tissue type including testis as equally relevant for calculating the PS of a given gene, as otherwise our list of CSGs would be exceedingly enriched in established cancer-testis antigens. However, as gender-specific normal tissue types such as uterus/ovary or prostate/testis, respectively, are irrelevant to nominate CSGs for the respective other gender, we calculated gender-specific CSG-scores omitting gender-specific tissue types for calculation of the PS of a given gene for the respective other gender (**Supplementary Table 1**). A meaningful CSG-score was determined statistically as being equal or above the 99.9^th^ percentile of all CSG-scores calculated across all cancer entities. Using this cut-off, the CSG-scores for CSGs potentially suitable for immunotherapeutic targets in a given cancer entity were usually greater than 2. CSG-scores greater than 3 were empirically considered as high and those greater than 4 as very high.

### Development of RAVEN (Rich Analysis of Variable gene Expressions in Numerous tissues)

We developed an application named RAVEN that incorporates several statistical methods to easily identify putative highly immunogenic peptides encoded by CSGs from any gene expression dataset. RAVEN and a detailed handbook as well as associated datasets can be downloaded free of charge and for academic use only under https://github.com/JSGerke/RAVENsoftware.

The graphical interface of RAVEN is simple and designed for scientists without bioinformatics background. The current program version developed with Java (for Windows, Linux and Mac) requires at least a Java 8 runtime environment.

RAVEN can interrogate gene expression datasets and compare expression levels of different genes in the same tissue or of the same gene in different tissues applying our algorithm as explained above. The statistical summary of such comparisons can be obtained in spreadsheet format and visualized by Java library JFreeChart. Additionally, the application enables users to retrieve either gene-or tissue-specific subsets of the interrogated gene expression dataset, which can then be further analyzed in RAVEN or other commonly used software such as Microsoft Excel or GraphPad Prism.

In addition, RAVEN includes a pipeline combining several bioinformatic services to offer a quick and simple way to obtain all peptide sequences of a pre-specified length (encoded by identified CSGs) and their affinity to different HLA-alleles. Furthermore, RAVEN nominates all MHCs that are predicted to present the identified peptides. To access the UniProtKB (UniProt Consortium, 2015) database via RAVEN, we used the UniProtJAPI library (Patient et al., 2008). To match gene IDs with their corresponding protein IDs of different databases such as UniProt and NCBI, RAVEN sends a query to the database identifier mapping service of UniProt. The implemented peptide-matching pipeline accesses the MHC-I binding prediction tool provided by the Immune Epitope Database (IEDB) Analysis Resource (Vita et al., 2015) via a RESTful interface (IEDB-API). T Cell Epitope Prediction identifies peptides binding to MHC class I of a certain protein sequence. Therefore, RAVEN uses artificial neural networks (ANN) and a prediction algorithm developed by NetMHC (Andreatta and Nielsen, 2016; Nielsen et al., 2003). The peptide search service (Chen et al., 2013) of UniProt is queried via a RESTful web service which API is provided and integrated by Protein Information Resource (PIR) using ApacheLucene for peptide text searches (Chen et al., 2013; Wu et al., 2003). In RAVEN, this approach is available for the most common alleles in human and mouse. In contrast to other methods provided by RAVEN, this pipeline is independent from the analyzed gene expression dataset but requires an internet connection.

### Human cell lines and cell culture conditions

Cells were grown at 37°C in humidified 5% CO_2_ atmosphere in RPMI 1640 medium (Biochrom, Berlin, Germany) supplemented with 10% FCS (Biochrom) and 100 U/ml penicillin and 100 μg/ml streptomycin (Biochrom). TAP-deficient HLA*A02:01^+^ T2 cell line (somatic cell hybrid) was obtained from P. Cresswell (Yale University School of Medicine, New Haven, CT, USA). T2 cells were maintained in RPMI 1640 medium additionally supplemented with 1 mM sodium pyruvate and non-essential amino acids (both Biochrom). Cell line purity was confirmed by short tandem repeat profiling (latest profiling 15^th^ December 2015) and cells were routinely examined by PCR for the absence of mycoplasma. A list of the used cell lines is provided in **Supplementary Table 2**.

### RNA extraction, reverse transcription and qRT-PCR

RNA was extracted with the Nucleospin RNA kit (Macherey-Nagel, Düren, Germany) containing a 15 min on-column DNA digestion step to degrade genomic DNA. RNA was reverse-transcribed using High-Capacity cDNA Reverse Transcription Kit (Applied Biosystems). qRT-PCRs were performed using SYBR Select Master Mix (Applied Biosystems). Oligonucleotides were purchased from MWG Eurofins Genomics (Ebersberg, Germany). Primer sequences are listed in **Supplementary Table 3**. Reactions were run in 10-20 μl final volume on a CFX Connect instrument and analyzed using the CFX Manager 3.1 (both Bio-Rad). Gene expression levels of specific genes were normalized to that of the housekeeping gene *RPLP0* (Martin, 2016).

### Human samples and ethics approval

Human tissue samples were collected at the Institute of Pathology of the LMU Munich (Germany) with approval of the corresponding institutional review boards. The ethics committee of the University Hospital of the LMU Munich approved the study (approval no. 307-16 UE).

### Immunohistochemistry (IHC) and evaluation of immunoreactivity

IHC analyses were performed on formalin-fixed, paraffin-embedded (FFPE) tissue samples. Paraffin blocks from several institutions were collected at the Institute of Pathology of the LMU Munich. From all blocks, we harvested 3 cores per sample with a core-diameter of 1 mm to assemble a tissue microarray (TMA). A list of the included tumor types and normal tissues is given in **Supplementary Table 4**. Of each TMA block 4 μm sections were cut and stained with an iView DAB detection kit (Ventana Medical System, Tucson, AZ) according to the company’s protocol. Subsequent antigen retrieval was carried out using TRIS-buffer and blockage of endogenous peroxidase with 7.5% aqueous H_2_O_2_. TMA sections were stained at a dilution of 1:180 for 60 min at room temperature with a monoclonal antibody against human PAX7 raised in mouse (Kawakami et al., 1997), which was purchased from the Developmental Studies Hybridoma Bank (Cat.No. PAX7-c; Iowa City, IA). Then slides were incubated with a secondary biotinylated anti-mouse IgG antibody (ImmPress Reagent Kit, Peroxidase-conjugated) followed by target detection using ABC-kit chromogen for 10 min (Dako, K3461).

At least three high-power fields (40x) of each core for every sample were assessed. Semi-quantitative evaluation of immunoreactivity was carried out by two independent physicians trained in histopathology. The intensity of immunoreactivity was determined as grade 0 = none, grade 1 = faint, grade 2 = moderate and grade 3 = strong. The particular percentage score of cells was assigned to 0, when any immunoreactivity was present in less than 20% of the cells, 20-39% to 1, 40-59% to 2, 60-79% to 3 and 80% or higher to 4. For calculation of overall immunoreactivity for the given protein, we multiplied the intensity score with the percentage score, in analogy to UICC guidelines for hormone receptor scoring in human breast cancer as described (Remmele and Stegner, 1987).

### Peptide binding assay using TAP deficient T2 cells

All peptides were solid-state synthesized with the highly-parallelized LIPS^®^ technology (Elephants & Peptides, Potsdam, Germany). As a positive control, we used an established highly affine influenza matrix protein epitope (M158-66; sequence GILGFVFTL) (Gotch et al., 1987). T2 cells were washed twice with PBS and seeded in round-bottom 96-well plates (TPP, Trasadingen, Switzerland) at a concentration of 2 × 10^5^ cells/well in a final volume of 200 μl. Cells were pulsed with increasing amounts of peptide to measure a concentration dependency of MHC-I binding. Unpulsed cells were used as a negative control. After incubation over-night, cells were washed twice with FACS-buffer consisting of PBS with 2% FCS and stained for HLA-A2 using a FITC mouse anti-human HLA-A2 antibody (BD Pharmingen™, Clone BB7) for 30 min at 4°C. For isotype control a BB515 mouse IgG2Ak antibody (BD Horizon™, Clone G155-178) was used. Then, cells were washed twice in FACS-buffer before being resuspended in PBS and analyzed using a FACSCalibur flow cytometer (Becton Dickinson). To determine the relative peptide binding, the fluorescence intensity of a peptide at a defined concentration was divided by the intensity of unpulsed T2 cells.

## RESULTS

### Dataset assembly, workflow, and basic concepts of RAVEN

In order to automatize the systematic and fast identification of CSGs as well as the prediction of corresponding highly affine peptides for any given MHC, we developed a user-friendly software named RAVEN (Rich Analysis of Variable gene Expressions in Numerous tissues). An overview on the workflow conducted by RAVEN is given in Fig. 1. The software, a detailed user guide and our gene expression datasets are freely available under https://github.com/JSGerke/RAVENsoftware.

**Figure 1.**
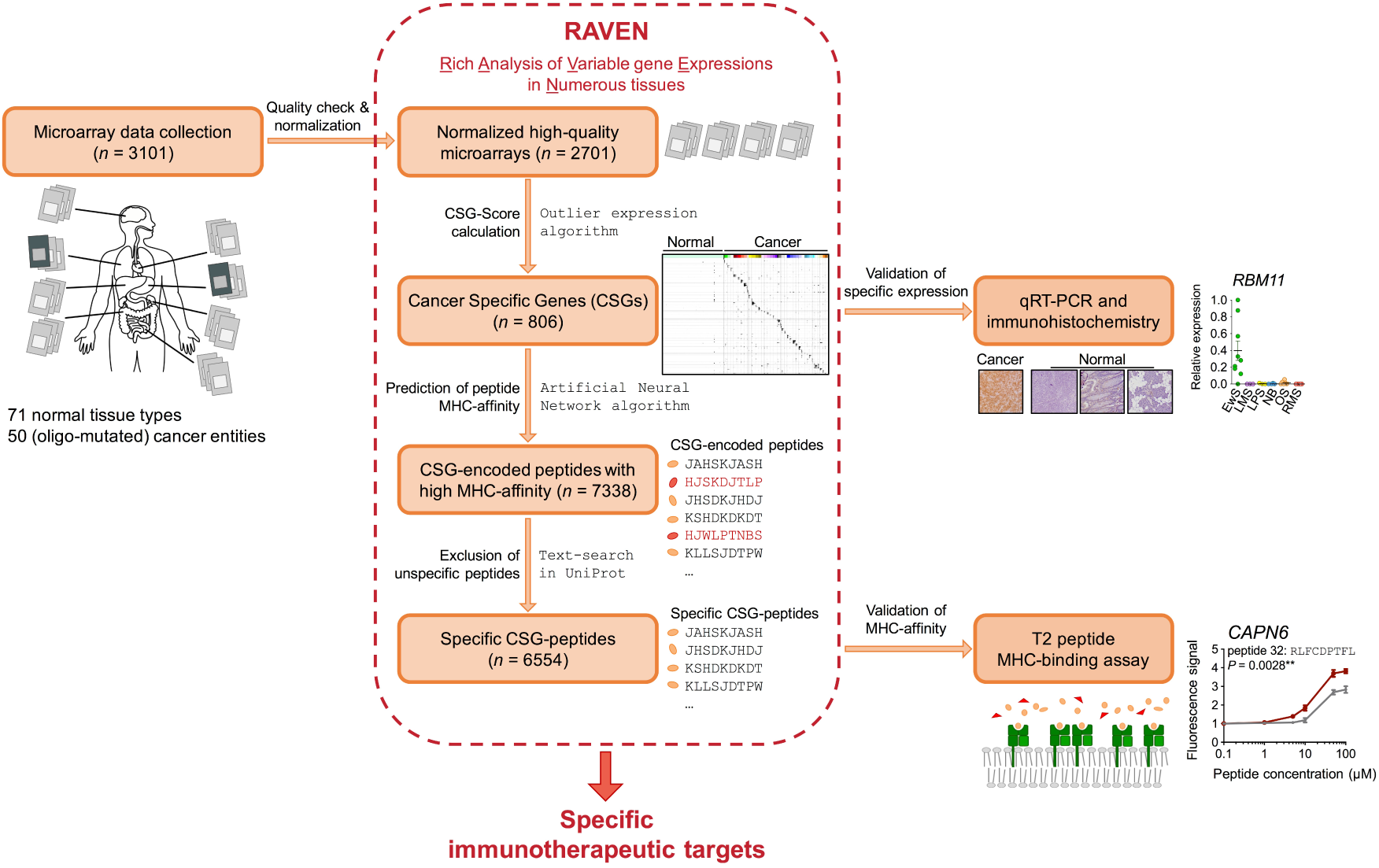
Schematic illustration of the assembly, quality-check, and normalization of gene expression data as well as tasks executed by RAVEN.

### Transcriptome-wide detection of CSGs overexpressed in multiple cancer entities with RAVEN

Previous studies have shown that many established CSGs are only expressed in subsets of specific cancer entities, which is often referred to as ‘outlier’ expression (Wu, 2007; Xu et al., 2012). Indeed, many CSGs are either virtually not expressed in somatic normal tissues or exclusively expressed in specific lineages such as embryonal and germline tissues (Monk and Holding, 2001; Simpson et al., 2005). This outlier expression discriminates cancer cells from normal somatic cells and may offer a therapeutic window for preferentially targeting cancer cells, e.g. by adoptive T cell therapy (Coulie et al., 2014). Also, it may increase the likelihood that lymphocytes responsive to the proteins encoded by CSGs are preserved in the mature lymphocyte repertoire (Simpson et al., 2005), because they are not counter-selected during lymphocyte development. However, an outlier expression profile implies that conventional statistical tests, which either simply aim at identifying generally upregulated CSGs across many cancer samples (e.g. Student’s t-test) or ignore the strength of overexpression in a small subset of patients (e.g. rank-based nonparametric tests), would fail to detect such clinically relevant CSGs.

Therefore, we developed a scoring algorithm to scan transcriptome-wide for CSGs by assigning an ‘outlier score’ (OS) to each gene for high expression in a given cancer entity, which is penalized by a ‘penalty score’ (PS) if high expression in any normal tissue type is present.

Both scores are calculated for each gene separately as the mean expression level of the 95^th^ and 75^th^ percentile. Then, we calculated an overall score for each gene named ‘CSG-score’, which is built by subtracting the gene-specific PS from the OS. This function highlights all genes overexpressed in only a subset of cancer samples, while avoiding the misrepresentation caused by extremely high outlier expression signals in single samples.

In addition, our algorithm takes into account gender-specific normal tissue types such as uterus and prostate. Specifically, our algorithm calculated gender-specific CSG-scores for each gene excluding normal tissues of sexual organs specific for the other gender (see Materials and Methods).

To analyze the expression profiles of human genes in normal and cancer tissues we compiled 85 Affymetrix HG-U133-Plus2.0 microarray datasets for 71 normal tissues and 50 cancer types with a focus on oligo-mutated pediatric cancers and sarcomas, totaling to 2701 high-quality and simultaneously normalized samples (**Supplementary Table 1**). In prospect of a future exploitation of our CSGs as clinical immunotargets, we included graft versus host disease (GvHD)-sensitive normal tissue types such as retina and colonic mucosa as well as normal B and T cells to obviate fratricide effects, which can compromise adoptive T cell therapies (Kirschner et al., 2016; Leisegang et al., 2010).

Applying our scoring algorithm to this well-curated gene expression dataset, RAVEN identified 806 non-redundant CSGs (defined by a CSG-score above the 99.9^th^ percentile of all scores across 50 cancer entities) (Fig. 2, **Supplementary Table 5**). Among them we found not only many established CSGs such as *LIPI* for Ewing sarcoma (Foell et al., 2008), *PRAME* for neuroblastoma (Oberthuer et al., 2004; Spel et al., 2015) and members of the *MAGE*-family for germinoma (Hara et al., 1999), neuroblastoma (Söling et al., 1999), synovial sarcoma (Iura et al., 2017), multiple myeloma (Condomines et al., 2007), diffuse large B cell lymphoma (DLBCL) (Hudolin et al., 2013), and osteosarcoma (Sudo et al., 1997), but also many novel candidates of which some appear to be suitable for targeting multiple cancer entities (Fig. 2, **Supplementary Table 5**).

**Figure 2.**
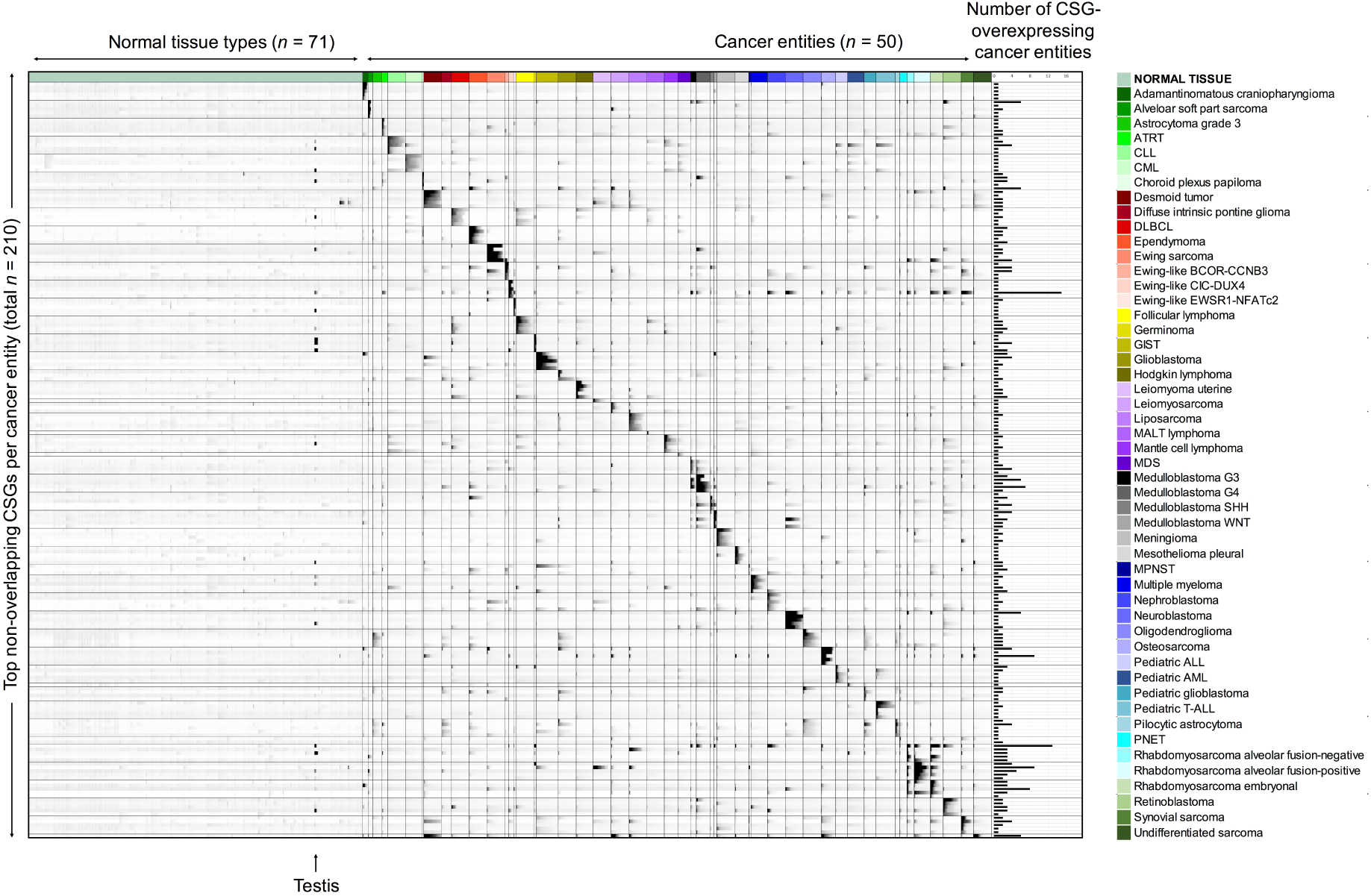
Overexpressed CSGs in multiple cancers entities identified with RAVEN. Relative gene expression intensities of the top-5 CSGs for each cancer entity excluding overlapping CSGs with other tumor entities re indicated in greyscale with black color representing high and white color low expression. Each line represents an individual CSG (for a complete list see **Supplementary Table 5**); each column represents one primary tumor/leukemia sample. The bar graph on the right displays the number of different cancer entities in which the corresponding CSG reached a CSG-score above the 99.9^th^ percentile of all CSG-scores. ALL, acute lymphoblastic leukemia; AML, acute myeloid leukemia; ATRT, atypical teratoid/rhabdoid tumor; CLL, chronic lymphatic leukemia; CML, chronic myeloid leukemia; DLBCL, diffuse large B cell lymphoma; GIST, gastrointestinal stromal tumor; MALT, mucosa associated lymphatic tissue; MPNST, malignant peripheral nerve sheath tumor; PNET, primitive neuroectodermal tumor.

The specific expression of nine selected CSGs was confirmed by qRT-PCR in a panel of cancer cell lines from six different tumor entities. As shown in Fig. 3A, there was a high concordance of calculated CSG-scores and expression intensities measured by microarrays in primary tumors with relative mRNA expression levels measured by qRT-PCR in corresponding cancer-derived cell lines.

**Figure 3.**
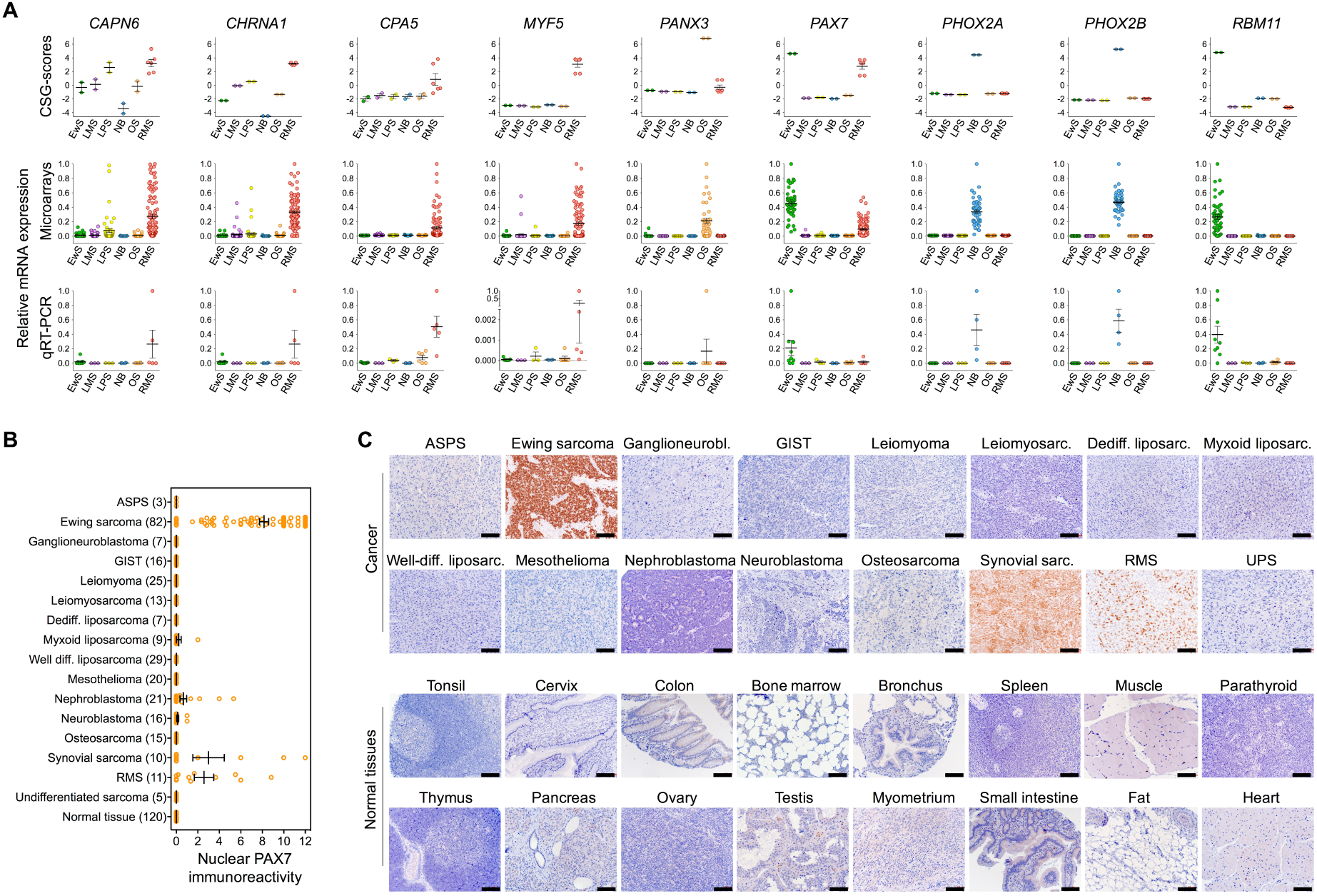
Validation of the expression pattern of selected CSGs by qRT-PCR and IHC. A)Upper and middle panel: CSG-scores and corresponding expression intensities (natural scale) of selected genes in primary Ewing sarcoma (EwS, *n* = 50), neuroblastoma (NB; *n* = 49), rhabdomyosarcoma (RMS; *n* = 101), liposarcoma (LPS; *n* = 50), leiomyosarcoma (LMS, *n* = 50) and osteosarcoma tumors (OS, *n* = 40). Lower panel: Relative expression levels of the same genes as determined by qRT-PCR in EwS (*n* = 9), NB (*n* = 4), RMS (*n* = 5) and LPS (*n* = 3), LMS (*n* = 3) and OS (*n* = 6) cell lines. B)Analysis of nuclear PAX7 immunoreactivity by IHC in indicated primary tumors and normal tissues. ASPS, alveolar soft part sarcoma; GIST, gastrointestinal stromal tumor. Numbers of analyzed samples are given in parentheses. *C)* Representative images of nuclear PAX7 IHC staining in cancer and selected normal tissues. Scale bar = 300 μm. UPS, undifferentiated pleomorphic sarcoma.

In particular, *PAX7* (paired box 7) showed a very high CSG-score (>5) in multiple cancer entities including oligo-mutated Ewing sarcoma. Therefore, we validated its strong overexpression on protein level in a subset of these cancer entities by immunohistochemistry in a comprehensive tissue microarray (TMA, *n* = 412 samples) also containing somatic and germline normal tissue types. As shown in Fig. 3B,C, *PAX7* was exclusively expressed in cancer entities with high CSG-scores, while being virtually not expressed in normal tissues. Collectively, these data demonstrate that RAVEN can reliably identify CSGs with specific overexpression in multiple cancers as compared to normal tissues.

**Figure 4.**
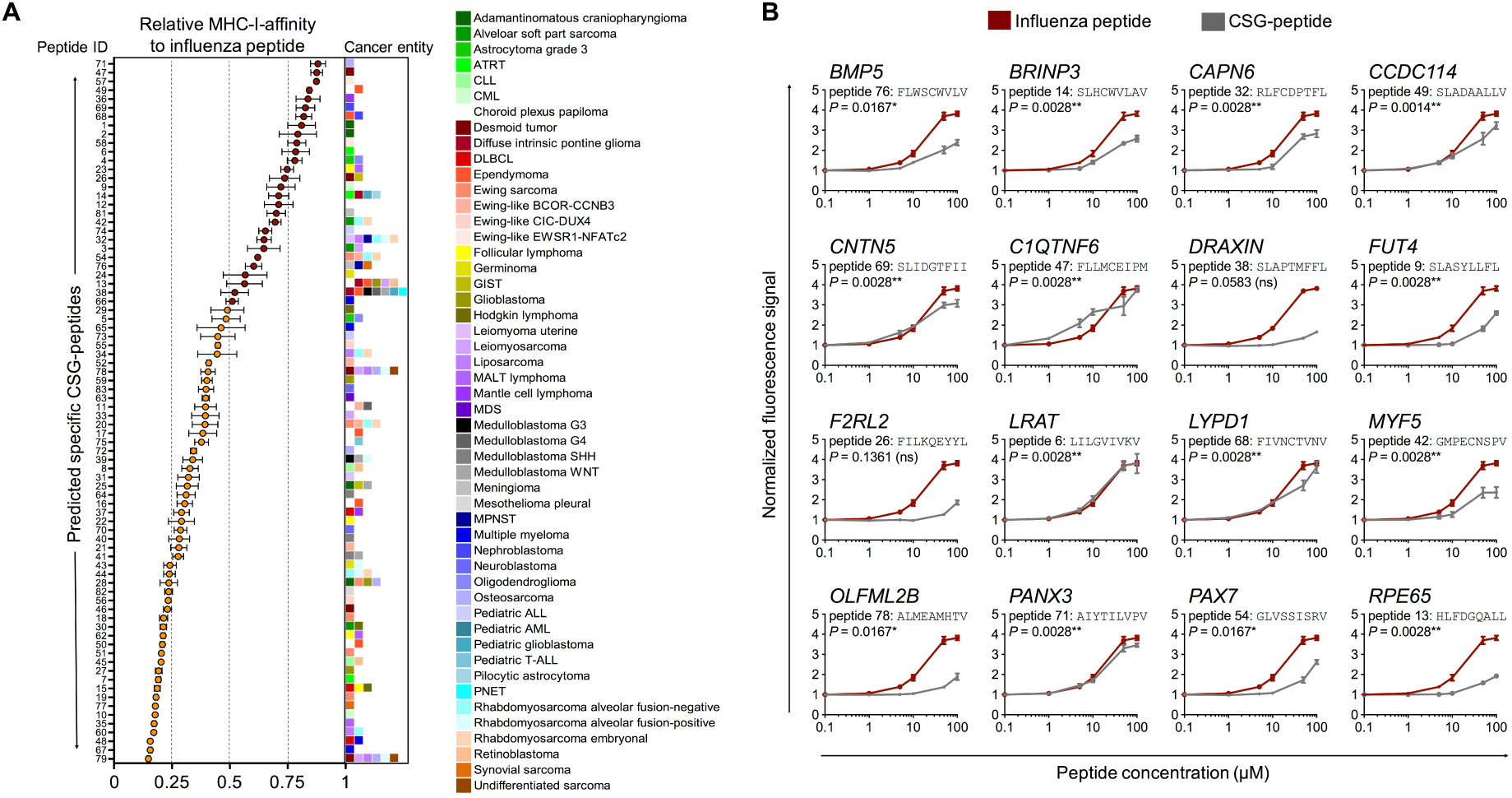
Validation of MHC-affinity of CSG-encoded peptides in a T2-binding assay. A)Relative MHC-I-affinity of 80 selected peptides at 100 μM in T2-binding assays as compared to a highly affine influenza peptide (peptide sequences are given in **Supplementary Table 6**). The colored boxes at the right side of the graph represent the number and type of cancer entities in which the corresponding CSG encoding the indicated peptide is overexpressed. Peptides with a MHC-affinity of ≥ 50% of the influenza peptide are highlighted in red color. Data are presented as mean and SEM of *n* ≥ 3 experiments. ALL, acute lymphoblastic leukemia; AML, acute myeloid leukemia; ATRT, atypical teratoid/rhabdoid tumor; CLL, chronic lymphatic leukemia; CML, chronic myeloid leukemia; DLBCL, diffuse large B cell lymphoma; GIST, gastrointestinal stromal tumor; MALT, mucosa associated lymphatic tissue; MPNST, malignant peripheral nerve sheath tumor; PNET, primitive neuroectodermal tumor. B)Normalized fluorescence signals of 16 selected peptides with high MHC-affinity as compared to that of a highly affine influenza peptide in T2-binding assays. Data are presented as mean and SEM of *n* ≥ 3 experiments. *P* values of a Spearman’s rank-order correlation are reported.

### Prediction of non-redundant CSG-encoded peptides with high MHC-affinity by RAVEN

To identify peptides encoded by CSGs suitable for a targeted immunotherapy, we implemented the artificial neural network (ANN) algorithm (Andreatta and Nielsen, 2016; Nielsen et al., 2003) provided by the immune epitope database IEDB 3.0 (Vita et al., 2015). RAVEN can apply this ANN algorithm to predict peptide-affinities for different peptide lengths and the most common human and murine MHC-subtypes.

In our list of 806 CSGs, RAVEN predicted potential highly affine peptides for 9-mers, which usually show optimal binding to most MHC class I molecules (Andreatta and Nielsen, 2016; Eichmann et al., 2014), and for HLA-A02:01, which is the most common MHC-I in Caucasians (González-Galarza et al., 2015). RAVEN automatically crosschecked these peptides by a text search algorithm with ApacheLucene (Chen et al., 2013; Wu et al., 2003) against the human reference-proteome (UniProt release 2015_06) to exclude sequence identity with non-specifically expressed proteins. In total, RAVEN predicted 7,338 9-mer peptides with high MHC-I-affinity (defined as a dissociation constant *K*_*d*_ ≤ 150 nM) of which 6,554 had no sequence identity with any other protein (**Supplementary Table 6**).

### Predicted CSG-encoded peptides exhibit strong affinity to MHCs

We next sought to confirm the predicted affinity of peptides to human HLA-A02:01 proposed by RAVEN. Therefore, we selected among the unique 6,554 peptides 80, which covered all 50 analyzed tumor entities and which had high to very high CSG-scores. For these 80 peptides, we designed a customized solid-phase synthesized peptide-library and assessed whether they can stabilize MHC-I on the surface of TAP2-deficient cells in T2-binding assays. As shown in Figure 4A, 38 of 80 tested peptides (47.5%) achieved at least 50% of the MHC-stabilizing effect of a highly immunogenic influenza control peptide (GILGFVFTL, **Supplementary Table 6**) at a saturation dose of 100 μM. For these CSG-peptides, we repeated the T2-assays with six different peptide concentrations (0.1 to 100 μM). Strikingly, some of them, including the one encoded by *PAX7*, showed MHC-stabilization kinetics similar to the influenza peptide (Figure 4B). Taken together, these experiments demonstrated that RAVEN can identify highly affine CSG-encoded peptides suitable for targeting multiple cancer types by leveraging publicly available gene expression data.

## DISCUSSION

High-throughput gene expression analyses of cancers and normal tissues generated comprehensive and freely available transcriptome datasets (Rung and Brazma, 2013). However, identification of CSGs and derivative peptides with high affinity to MHCs continued to be laborious and slow (Gubin et al., 2015).

Here, we reported on the development and application of a mathematical scheme for transcriptome-wide detection of CSGs and their corresponding highly MHC-affine peptides as immunologic and clinical targets, and provide a use-friendly software (RAVEN), which automatizes this process. Applying RAVEN to a large gene expression dataset comprising multiple and often oligo-mutated pediatric cancer types as well as a broad spectrum of normal tissues revealed many CSGs with diagnostic and therapeutic potential. Moreover, we provide an analogous dataset including 19 of the most common carcinoma entities (1,462 samples; **Supplementary Table 1**, https://github.com/JSGerke/RAVENsoftware), which can be used for CSG identification in these tumor types.

In our pediatric cancer dataset, we observed significant enrichment (*P* < 0.0001, Chi^2^-test) of established cancer-testis antigens (CTDatabase, www.cta.lncc.br, (Almeida et al., 2009)), but also identified many novel candidates including *PAX7*. *PAX7* encodes a paired box transcription factor required for embryonal neural development (Kawakami et al., 1997) and renewal of skeletal muscle stem cells (Oustanina et al., 2004). Translocations involving *PAX7* and *FKHR* are found in the majority of alveolar rhabdomyosarcomas (ARMS), indicating a role of *PAX7* in the pathogenesis of myogenic tumors (Barr, 1999). Using RAVEN, we identified *PAX7* as a strong CSG in multiple oligo-mutated cancer entities such as Ewing sarcoma, Ewing-like sarcomas with a BCOR-CCNB3-translocation and embryonal as well as alveolar fusion-negative rhabdomyosarcoma. Its exclusive expression in these cancer entities was confirmed on protein level by IHC. Strikingly, *PAX7* encodes a 9-mer peptide (GLVSSISRV) with very high affinity for the most frequent MHC-I subtype in Caucasians (HLA-A02:01) (González-Galarza et al., 2015), rendering *PAX7* as an attractive target for immunotherapy for multiple oligo-mutated cancers.

The parameters of the analysis applied in RAVEN have been optimized to discover CSGs, which are virtually not expressed in most somatic tissues. Although some identified CSGs did not encode peptides suitable for immunotargets, a subset of them could constitute interesting targets for conventional pharmacotherapy. In fact, the CSGs *FGFR4*, *CDK4*, and several *MMPs*, which are specifically overexpressed in rhabdomyosarcoma (*FGFR4*), liposarcoma (*CDK4*), and desmoid tumors, leiomyoma, osteosarcoma and adamantinomatous craniopharyngioma (*MMPs*) (**Supplementary Table 5**), respectively, could be targeted by specific inhibitors currently in clinical trials (Chen et al., 2017; Hagel et al., 2015; Matziari et al., 2007).

Besides their potential utility as (immune)-therapeutic targets, some CSGs may harbor the potential to serve as diagnostic markers: While CSGs expressed in multiple tumor entities could be utilized for cancer-screening, CSGs exclusively expressed in certain cancer types can be used to identify and differentiate specific tumor entities. This could be important for determining treatment options, which is often difficult in cancers of unknown primary.

As RAVEN can also be applied to datasets only containing cancer samples, RAVEN can easily identify potential diagnostic markers among several cancers in parallel. In principle, our work-flow embedded in RAVEN provides an unbiased approach for transcriptome-wide detection of CSGs, which can be adapted to many specific applications, such as the identification of autoantibody signatures, biomarkers, tumor vaccine targets, or membrane antigen targets. Its performance could be further enhanced by combining it with other datasets, on cancer plasma or membrane proteomics. Since our algorithm provides a quantitative and gender-specific value for each gene in each tumor entity (**Supplementary Table 5**), the preferential expression of each CSG in different cancers is apparent at a glance. With more and more deep transcriptome sequencing data available and the advent of digital gene expression technology, we expect that RAVEN will be a highly beneficial tool to maximize the identification of CSGs and, hence, new diagnostic markers and therapeutic targets based on these data.

## ACKNOWLEDGEMENTS

We thank Mrs. Andrea Sendelhofert, Mrs. Anja Heier and Ms. Mona Melz for excellent technical assistance.

## AUTHOR CONTRIBUTIONS

MCB, JSG and TGPG conceived the study, performed bioinformatic and wet-lab analyses, and drafted and wrote the paper. MFO helped with bioinformatic analyses and assembly of gene expression datasets. ME, KS, and DHB provided immunological guidance and helped in experiments. MMK, NA, ANA, FCR, ÖÖ, TK, SS, TH, HS and DB contributed to the tissue microarray. JSG programmed and developed RAVEN. AK and UT performed T2-cell assays. FB and TF provided immunological guidance. MD, RAR, JM, AM, SO, MMLK, GS helped in wet-lab analyses. JL helped in IHC experiments. TK provided laboratory infrastructure, tissue samples and histological guidance. TGPG supervised the study and performed histological analyses. All authors read and approved the final manuscript.

